# *Streptomyces coelicolor*-plant association facilitates ergothioneine (EGT) uptake in *Triticum aestivum*

**DOI:** 10.1101/2025.04.08.647769

**Authors:** Alexandra Pipinos, Jinjun Kan, Andrew Smith, Gladis Zinati, Harsh Bais

## Abstract

The growing market of agricultural biologicals as alternatives to synthetic crop chemicals is driven by their ability to improve soil health, reduce carbon footprints, enhance crop yield and quality, and help counter declining protein levels in cereal crops linked to climate change and soil degradation. Ergothioneine (EGT), an amino acid with recognized nutraceutical and micronutrient properties, has gained popularity for its anti-inflammatory and antimicrobial properties on human health. While plants and humans cannot biosynthesize EGT, its production by *Streptomyces coelicolor* presents as a promising bio-stimulant to support overall plant health and human health. Our study investigates the potential for *Streptomyces coelicolor* M145 to enhance EGT levels in spring wheat (*Triticum aestivum*). Results confirmed successful EGT extraction from bacterial cell extracts and plant tissues. Additionally, a fluorescent confocal microscopy staining and imaging protocol showed bacterial colonization on *T. aestivum* and its potential as a root endophyte. Following root inoculation, *S. coelicolor* was observed inhabiting roots, shoots, and internodes of *T. aestivum*, suggesting its potential endophytic lifestyle on host plants. Our data showed that *S. coelicolor*-associated wheat plants produce EGT in planta. Overall, our findings establish a direct link between soil and human health through rhizosphere colonization by *S. coelicolor* and in planta production of EGT, suggesting an alternate route to enhance protein concentration in crop plants.

## Introduction

The connection between plant nutrition and human health is a critical area of research, especially as the global population is projected to exceed 10 billion by 2050^1^. To address the growing demand for nutrient-rich food, researchers are focusing on fortifying the world’s most widely cultivated crops—such as wheat, soybean, corn, and rice—with essential nutrients and bioactive compounds^2^.

Compared to synthetic fertilizers, agricultural biologics, introduced in the mid-20^th^ Century, are natural substances or biological organisms that enhance overall plant health and growth^3^. With a projected growth to reach around $24.6 billion by 2027, agricultural biologics or microbial inoculum synthetic communities (SynCom) are emerging as a promising alternative to synthetic treatments, while synthetics have come under scrutiny due to their environmental and health impacts ^4^. Agriculture accounts for 12% of global greenhouse gas (GHG) emissions, with a significant portion attributed to dinitrogen monoxide released from nitrogen compounds in chemical fertilizers^5^. Moreover, synthetic inputs in crop production may pose risks to human health, having associations with an increased likelihood of certain disease development, including increased risk of asthma, chronic obstructive pulmonary disease (COPD), and lung cancer ^5,6^. In fact, the EU has introduced policy measures through the Farm to Fork strategy, established in 2020. This strategy aims to achieve targets such as a 50% reduction in pesticide use and risk, in addition to 20% reduction in chemical fertilizer use by 2030^7^. With chemical-based becoming less frequently used, agricultural biologics offer a more sustainable approach to enhancing crop nutrition while minimizing harmful effects on the environment and human health ^3^. Agricultural biologics are often categorized into three main subgroups: biofertilizers, biopesticides, and bio stimulants^8^.

Plant bio stimulants are substances or microorganisms that can be added to plants to increase nutrient efficiency, abiotic stress tolerance, crop quality traits, and other beneficial impacts to overall crop health^9^. Zhang and Schmidt explored the use of humic acids and seaweed extracts, demonstrating their potential to promote plant growth^10^. In addition to seaweed extract, which remains the most extensively studied component of bio stimulants, other substances that can function as bio stimulants include amino acids, microbials, plant extracts, and organic acids^11^. Key microbial bio stimulants include Plant-Growth Promoting Rhizobacteria (PGPRs), which, in many cases, positively impact plant health with or without directly providing nutrients^9^.

Actinobacteria, a diverse phylum of soil-inhabiting, Gram-positive, filamentous bacteria that support nitrogen fixation, phosphate solubilization, iron acquisition, and overall crop development ^12^. Within this group, *Streptomyces* is especially notable for its antimicrobial properties and its production of bioactive compounds that enhance plant growth^13,14^. As natural soil dwellers, *Streptomyces* colonize the rhizosphere both externally and endophytically, improving soil fertility and nutrient uptake^15,16^ Interest in these microbes has grown as declining crude protein levels in cereal crops – driven by climate change, plant breeding, declines in soil organic matter, and changes in environmental legislation – have raised concerns about malnutrition and nutrient deficiencies in vulnerable populations^17^ contributed to a decline in crude protein content in agricultural crops^171819^. Therefore, exploring methods to enhance protein levels in common cereal crops has become a critical focus in agricultural research.

Ergothioneine (EGT) is a unique, naturally occurring amino acid with potent antioxidant and anti-inflammatory properties that are beneficial to human health^20^. Examples of EGT’s potential linkages in health benefits include its association with reduced risk of cardiovascular disease and mortality^21^.

Additionally, increased EGT consumption has been linked to healthy cognitive aging ^22^. EGT also exhibits protective qualities for the skin, including improving skin hydration, enhancing elasticity, and reducing trans-epidermal water loss^10^. These benefits highlight its increasing popularity in skincare research. Although EGT offers significant benefits to human health, it is neither produced in humans nor plants ^23^. As a result, EGT must be obtained through diet, either from food sources or in supplement form. While dietary sources of EGT are primarily limited to mushrooms, a few fungi and bacteria have adapted the ability to biosynthesize it. Notably, Actinomycetes, cyanobacteria, and methylobacteria are the only known bacteria class capable of biosynthesizing EGT ^20^. Some fungal species such as *Neurospora crassa, Cordyceps militaris*, and mushroom fruiting bodies also produce ERGO ^24^. In particular, a member of the Actinobacteria class, *Streptomyces coelicolor* M145 contains the five-gene biosynthetic cluster to biosynthesize EGT ^25^. Therefore, considering *S. coelicolor*’s role as a PGPR and its ability to produce EGT, using the organism as a bio stimulant to enhance EGT levels in a popular cereal crop-compensating for protein loss- is of high interest.

To advance our understanding of EGT production in *Streptomyces coelicolor* M145 and its potential as a bio stimulant for enhancing crop health and nutrition, we conducted controlled experiments to evaluate *S. coelicolor* both independently and in conjunction with the inoculation of a widely consumed cereal crop, spring wheat (*Triticum aestivum*). A method was developed for the detection and quantification of EGT using Ultra-Performance Liquid Chromatography (UPLC) coupled with Tandem Mass Spectrometry analysis. Our data showed an effective protocol for extracting and quantifying EGT from intracellular *S. coelicolor*, comparing intracellular EGT levels at days 5, 7, and 10 post-inoculations. Via a co-inoculation experiment, we further showed that *S. coelicolor* robustly binds with wheat roots.

This was demonstrated by developing a protocol for fixation, staining, and fluorescent confocal imaging to qualitatively assess the interactions between *S. coelicolor* and *T. aestivum.* Our data also showed that presence of EGT in plants post-inoculated with *S. coelicolor* at various time points. Through these experiments, we demonstrate that *S. coelicolor* functions as a biostimulant, enhancing EGT concentration in *T. aestivum* plants. Our findings support the use of crop biologicals as a viable strategy to increase overall protein concentration, addressing the growing concern of protein deficiency in staple crops.

### Materials and Methods Chemicals

L-Ergothioneine (L-EGT, purity > 99.99%) was obtained from MedChemExpress (Monmouth Junction, NJ). Ammonium acetate, methanol (For HPLC, >=99.9%), water (HPLC grade), and acetonitrile were purchased from Sigma-Aldrich Inc (St. Louis, MO). For microscopy staining, Wheat Germ Agglutinin (WGA) 594 was obtained from Thermo Fisher Scientific (Waltham, MA) and the calcofluor white stain was obtained from Sigma-Aldrich Inc (St. Louis, MO).

### Instruments

The Xevo-G2-S QTof system by Waters^TM^ (Milford, Massachusetts) was utilized for Ultra-Performance Liquid Chromatography (UPLC) of EGT sample analysis. The Andor Dragonfly confocal super resolution microscope by Oxford Instruments (Abingdon, United Kingdom) was used in the fluorescent confocal microscopy analysis.

### Plant material

*Triticum aestivum* Surpass Spring seeds were obtained from South Dakota State University (Brookings, SD), and were stored long-term in the dark at room temperature inside their original cloth bag. The seeds were sanitized prior to plating, and the husks were removed by light force. Using sterilized forceps, wheat seeds were transferred to petri plates containing 25 mL of full-strength Murashige and Skoog^26^ media per plate. The plates were sealed with micropore tape or parafilm and incubated under light at 26°C for 7 days to germinate.

### Bacterial Culture

*Streptomyces coelicolor* M145 strains were donated by Matthew Traxler lab at the University of California Berkeley. Using a sterile inoculation loop, the culture was streaked onto a solid, soy-flour mannitol (SFM) media plate. The *S. coelicolor* plates were sealed with micropore tape and incubated at 30° C for 3-5 days until gray-white colonies appeared. For longer-term storage, a 20% glycerol stock was prepared from the original plates and stored in a -80° C freezer.

### Preparation of Standard Solution and Standard Curve

To create a 10 mM stock solution of L-EGT, 2.293 mg of L-EGT was dissolved in 1 mL of 50% methanol extraction solvent. The stock solution was serial diluted using the extraction solvent to prepare standard solutions at a concentration range of 0.01-10 µM. These standards were used to generate a calibration curve. EGT presence was confirmed using MS/MS tandem mass spectrometry to confirm precursor ion along with product ion presence (SOM Fig. 1). To account for variations in sample matrices, separate standard curves were generated to determine an appropriate limit of quantification (LOQ) for all sample extract experiments.

**Figure 1:**
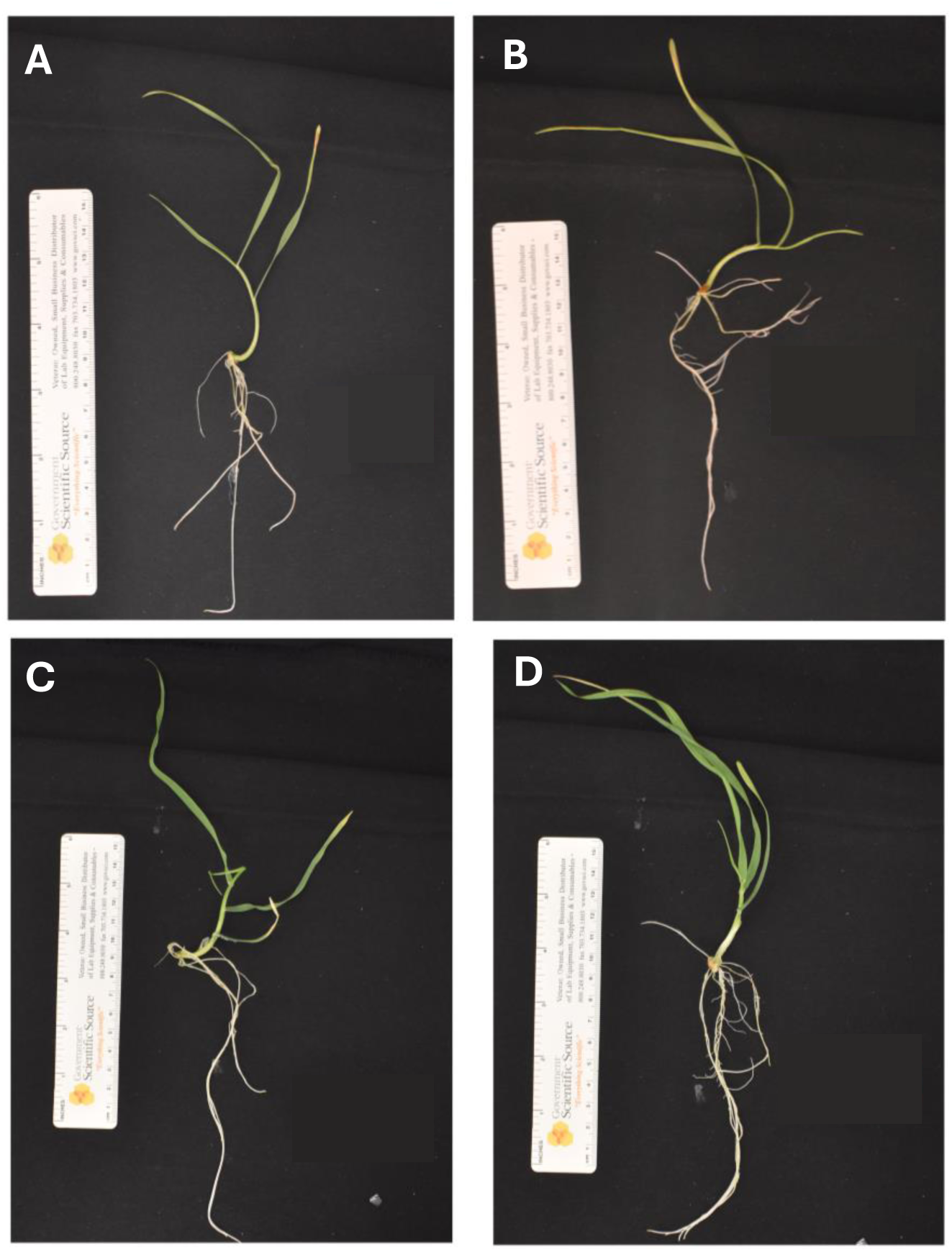
Representative images of plants at day 5 and 10 post-inoculation on the day of harvest, the panels (A-B) showed control and *S. coelicolor* inoculated plants on day 5. The panels (C-D) shows control and *S. coelicolor* inoculated plants on day 10. Scale bar= 1.0 cm.

### Intracellular Cell-Extraction from *Streptomyces coelicolor* M145 and MS/MS Tandem Mass Spectrometry Quantification

Bacterial cells were harvested for EGT quantification on days 5, 7, & 10 of incubation. Approximately 10 mg of the bacterial inoculum was removed from the culture plate and transferred to a centrifuge tube containing 10% acetonitrile. The sample was vortexed for 10 seconds, then sonicated for 10-15 minutes. After sonication, the sample was transferred to a centrifuge (5430/5430R centrifuge, Eppendorf, Hamburg, Germany) at 4° C for 10 minutes at 12 000 x g. The supernatant was then transferred to a falcon tube. To ensure all EGT was collected, 1,000 µL of 10% acetonitrile was added to the centrifuge tubes and re-centrifuged under the same conditions and the extra supernatant was collected. The samples were lyophilized for two days and were then dissolved in 50% methanol. The samples were filtered, and a spiking solution was added to each sample. The samples were then subjected to Ultra-Performance Liquid Chromatography tandem mass spectrometry analysis (UPLC-MS/MS).

### *Streptomyces coelicolor* M145 inoculation on Surpass Spring Wheat

For the inoculation and colonization experiment, *T. aestivum* seeds were sterilized and plated on full strength Murashige & Skoog (MS) medium. The seeds were left to germinate under a 12-hour photoperiod at 26° C and 25% humidity for 7 days. After the spring wheat seeds were germinated by day 7, the plants were transferred to 10% MS and were inoculated with 10^6^ CFU/mL of *S. coelicolor* suspension. Control plants were incubated with a bacteria-free suspension.

### Ergothioneine Extraction from Inoculated Surpass Spring Wheat

For the EGT extraction of plants, a root or shoot sample was ground to a powder using liquid nitrogen and transferred to a tube containing 10% acetonitrile. The samples were briefly vortexed, sonicated for 10 minutes, then centrifuged at 4° C for 10 minutes at 12 000 x g. Once centrifugation was complete, the supernatant was transferred to a falcon tube. The pellet was re-extracted with more 10% acetonitrile, vortexed, centrifuged under the same conditions, and the supernatant was collected. The samples were then lyophilized for 48-72 hours. The lyophilized product was then dissolved in 50% methanol, filtered, and subjected to Ultra-Performance Liquid Chromatography tandem mass spectrometry analysis (UPLC-MS/MS).

### EGT detection by UPLC-MS/MS & Overall Protein Concentration Assay

EGT detection was carried out on a Waters Xevo G2-S QTof, with an electrospray ionization probe operated in positive mode. A custom protocol was developed in-house for EGT detection and quantification. Compound separation was conducted on an Intrada Amino Acid column, (50 mm x 3.0 mm, standard pressure) with (solvent A)- acetonitrile containing 0.1% formic acid, and (solvent B)- 100mM ammonium acetate. The flow rate for solvent A and solvent B were run at a consistent 0.5 mL/min. Solvent A was run at 95% from initial time until 4 minutes, then reduced to 5% between 4-4.5 minutes. At 4.5 minutes, solvent A was increased to 95% until the end of the run, to 6 minutes. Solvent B was run at 5% from the initial time to 4 minutes, then adjusted to 95% from 4-4.5 minutes. At 4.5 minutes, solvent B was reduced to 5% until 6 minutes. The autosampler was maintained at 4.0° C, and the column temperature was managed at 40°C. EGT analysis was directed using the Waters MassLynx software (version 4.2), where EGT peaks were analyzed and the area under the curves were integrated to determin the concentration of EGT. **A** Bradford assay using the BioRad Quick Start™ Bradford Protein Assay (Item #: 5000201) was performed to measure protein levels in plant roots and shoots. Spring wheat plants inoculated with ∼10^6^ cells/mL of *S. coelicolor*, along with control plants, were harvested on day 10 post-inoculation. The same protocol for EGT extraction from plants was used for the protein extraction in this assay. The OD_595_ measurements were taken using the Bio-Rad SmartSpec 3000 UV/VIS Spectrophotometer, and the linear regression equation generated from the standard curve was used to calculate the concentration (mg/mL) of each unknown sample.

### Fixation, staining, and confocal fluorescence microscopy of plants colonized by *S. coelicolor*

*Triticum aestivum* plant samples were sectioned into distinct regions of the plant and transferred to a 24-well plate containing a fixative composed of 25% glutaraldehyde, 4% paraformaldehyde, 100% Triton X-100, and 1x phosphate buffer solution (PBS). The samples then underwent vacuum filtration for one hour at 4° C, and the fixative was then removed. Samples were rinsed with phosphate buffer solution (PBS), and 0.2 M glycine was added into each well, where the samples were placed in a fridge (4°C) overnight. Plant samples were again washed three times with PBS and stained using Calcofluor white and Wheat Germ Agglutinin 594 (WGA594). Finally, solution Scale P (6 m urea, 30% (v/v) glycerol, and 0.1% (v/v) Triton X-100 in 1X PBS pH 7.4) was introduced into each well. The samples were stored at room temperature in a dark room until fluorescent confocal microscopy imaging. Z-stacking was performed for each region of the plant sample displaying colonization of bacteria using the Leica DMi8 widefield scope as a base coupled with the Andor Dragonfly 600 Spinning Disc Confocal imaging scope located in the University of Delaware Bioimaging center. Calcofluor white was excited using a 405 nm laser with emission captured at 446 nm, and Alexa fluor 594 was excited at 561 nm with emission captured at 594 nm. Z stacks were analyzed using Imaris Microscopy Image Analysis Software by Oxford Instruments.

### Endophyte Assay

Treated *Triticum aestivum* shoots and internodes along with untreated control plants were sectioned into various regions using a sterile scalpel and forceps. The cut plant samples were briefly immersed in 50% ethanol for 15 seconds, rinsed with sterile water, and placed onto soy-flour mannitol (SFM) plate. Samples were incubated at 30° for 3-5 days and subsequently examined for *S. coelicolor* growth.

### Statistical Analysis

The data was analyzed using GraphPad Prism (GraphPad Prism, version 10.4.0, Boston, MA). The following tests were performed for data analysis: One-way ANOVA statistical test paired with the Brown-Forsythe and Welch test, Dunnett’s T3 multiple comparisons test, Welch’s test, all with a significance level of P < 0.05.

## Results

### Inoculation of *Streptomyces coelicolor* M145 on *Triticum aestivum* enhances overall root and shoot fresh weight

Representative images of the harvested plants are shown in Fig. 1. A significant increase in root and shoot biomass was observed 10 days post-inoculation with *Streptomyces coelicolor* M145 (See SOM Fig. 2). These results suggest that *S. coelicolor* acts as a plant growth-promoting rhizobacterium (PGPR) by enhancing *T. aestivum* fresh weight in both roots and shoots.

**Figure 2:**
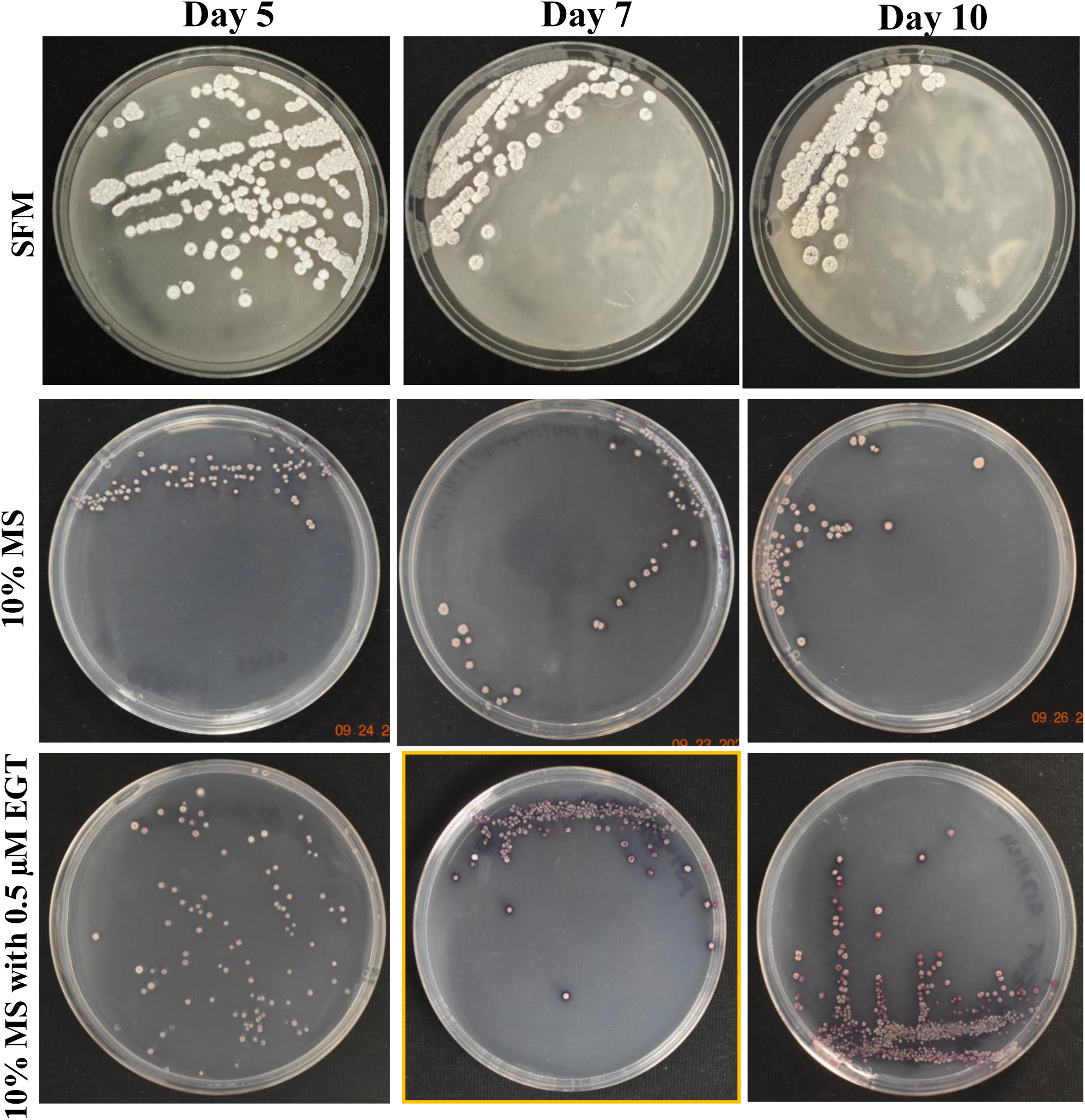
Growth of *Streptomyces coelicolor* on day 5, 7, and 10 across three media types. Streptomyces coelicolr was plated on soy-flour mannitol (SFM), 10% Murashige & Skoog (MS), and 10% MS supplemented with 0.5 μM L-EGT. The images illustrate morphological changes and growth patterns under varying nutritional conditions.

### Exogenous EGT supplementation induces morphological changes in *Streptomyces coelicolor* M145

Growth of *Streptomyces coelicolor* M145 varied across varying media conditions, with morphological characteristics including pigmentation being documented (Fig. 2). When *S. coelicolor* was grown on its preferred rich media, SFM, morphological characteristics of the cells were relatively consistent across days 5, 7, and 10 of growth. The colonies were relatively larger, with clear and consistent mycelial formations visible in each colony. The colonies appear gray/white throughout all three time points, with one noticeable feature on day 10 where one-two colonies are shown to be producing actinorhodin (purple color), a secondary metabolite that is oftentimes produced when *S. coelicolor* is under nutrient-limiting or stressful conditions. When *S. coelicolor* was grown under limiting condition with 10% MS, the size of the colonies averages a smaller diameter compared to the SFM grown cells, with inconsistency in diameter (See Fig. 2). The recorded colonies were still gray/white in color, but there was a lack of clear mycelial growth compared to the SFM cells (see Fig. 2). The production of actinorhodin was seen to progress with inoculation time under limiting growth conditions (Fig. 2). To evaluate the role of exogenous EGT, *S. coelicolor* cells grown on 10% MS media were supplemented with 0.5 µM L-EGT. On day 5, for *S. coelicolor* cells have signs of actinorhodin production both intracellular and extracellularly (see Fig. 2). Similarly to the 10% MS cells, there is inconsistent diameter size of the bacterial colonies in the 10% MS with L-EGT. Some of the colonies appear gray/white. By day 7 of growth, the cells appear to have produced significantly higher levels of actinorhodin compared to day 5. The day 10 image appears similarly to day 7, with most of the colony’s purple from actinorhodin synthesis, relatively small colony diameters compared to SFM and 10% MS cells, and a lack of clear morphological development such as mycelial formations. Overall, it can be interpreted that providing exogenous 0.5 µM L-EGT to a minimal media induces elevated stress to *S. coelicolor* cells by the visually enhanced production of actinorhodin.

### EGT production in *Streptomyces coelicolor* M145 grown under different media conditions

Growth of *Streptomyces coelicolor* varied across different media conditions. On day 5 of harvest, SFM and 10% MS cells produced very similar levels of EGT, where cells grown on 10% MS averaged insignificantly lower concentrations of EGT compared to SFM (See Fig. 3). By day 7, there was a statistically significant difference in EGT production comparing the two media conditions. *S. coelicolor* cells grown on SFM produced an average of ∼0.32 µM of EGT, while 10% MS cells were producing an average of 0.27 µM (See Fig. 3). The same level of statistical significance is shown comparing day 10 under both the media conditions (SFM and 10% MS), where *S. coelicolor* cells grown on SFM revealed higher EGT production compared to 10% MS (average ∼0.31 µM SFM, ∼ 0.27 µM 10% MS). The data suggests that EGT production in *S. coelicolor* is modulated with the culture period and media conditions. The data also reveals that a rich media condition may facilitate more EGT production in *S. coelicolor* cells.

**Figure 3:**
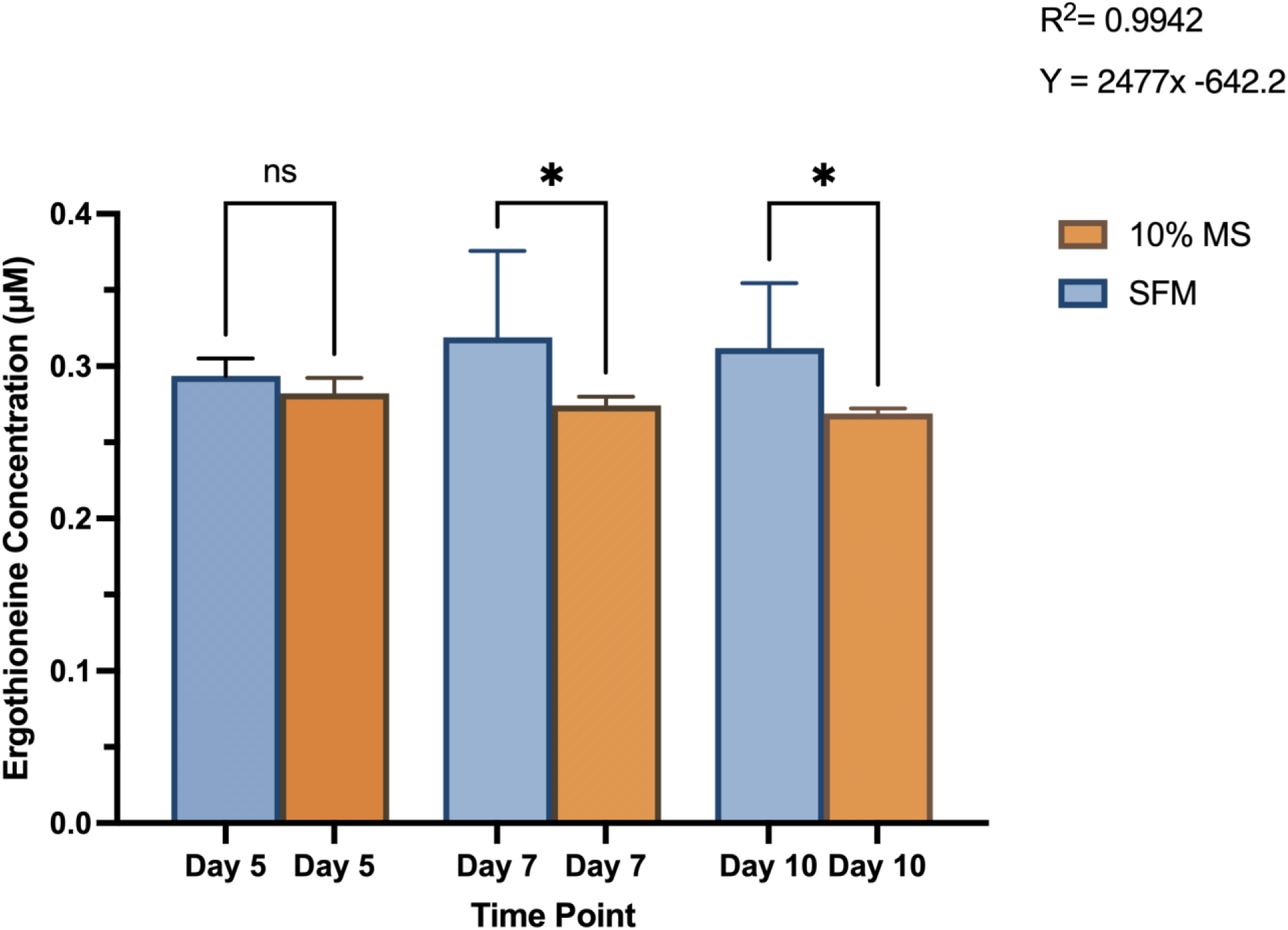
Intracellular ergothioneine (EGT) levels in *Streptomyces coelicolor* M145 cells grown on a plant growth medium, 10% Murashige & Skoog (MS) and Soy-Flour Mannitol (SFM) on days 5, 7, and 10 of growth. EGT levels are displayed in μM. Asterisks indicate statistically significant differences (p < 0.05) between treatments.

### Bioimaging reveals different lifestyles of *Streptomyces coelicolor* M145 on plant host *Triticum aestivum*

We showed that *S. coelicolor* biosynthesizes EGT and inoculation of *S. coelicolor* on host plants (wheat) promotes growth, next we evaluated the colonization patterns of *S. coelicolor* on wheat plants using live imaging. *Triticum aestivum* plants inoculated with ∼10^6^ cells/mL of *S. coelicolor* were harvested for confocal fluorescence microscopy on different days post inoculation (Figs. 4-6). The control untreated plants were imaged to reveal the gnotobiotic nature of the experimental condition devoid of any contaminants (see SOM Fig. 3). To monitor the temporal colonization pattern of *S. coelicolor* on wheat polants, we investigated different area of colonization throughout the wheat plants. The regions of interests for colonization are depicted as schematic in Fig. 4. We also developed a staining protocol using a fluorescent dye WGA594 for *S. coelicolor*. The regions of interest were divided into five different plant compartments [root tip (a), lateral root (b), crown (c), node (d) and leaf blade (e)] to evaluate the presence of *S. coelicolor* on wheat plants. Plants were imaged days 5/7 and 10 post inoculation with *S. coelicolor* and different regions were compared for the presence of *S. coelicolor*. As early as day 5 post inoculation, *S. coelicolor* was observed in the root tip and elongation zone, with threads of vegetative hyphae clearly visible and appear to be linking on to a root hair (Fig. 4). Temporally, *S. coelicolor* was observed to be present and associated with the lateral root regions mainly colonizing the root hairs post day 5 of inoculation. Like the root tip region, there are evident vegetative and aerial hyphae formations. However, in this region, the structural formations of *S. coelicolor* are more extensive as compared to the root tip regions (Fig. 4), where a larger colony is seen colonizing the root hairs. The crown region shows more thorough distribution of *S. coelicolor* with visible vegetative mycelia clasping the mature root hairs (see Fig. 4). The node region post day 5 of inculation showed presence of *S. coelicolor* colonizing the trichomes (see Fig. 4). Interestingly, when the leaf blades were analyzed post fixing and staining, apotential colonization by *S. coelicolor* was observed in the cut regions (See Fig. 4).

**Figure 4:**
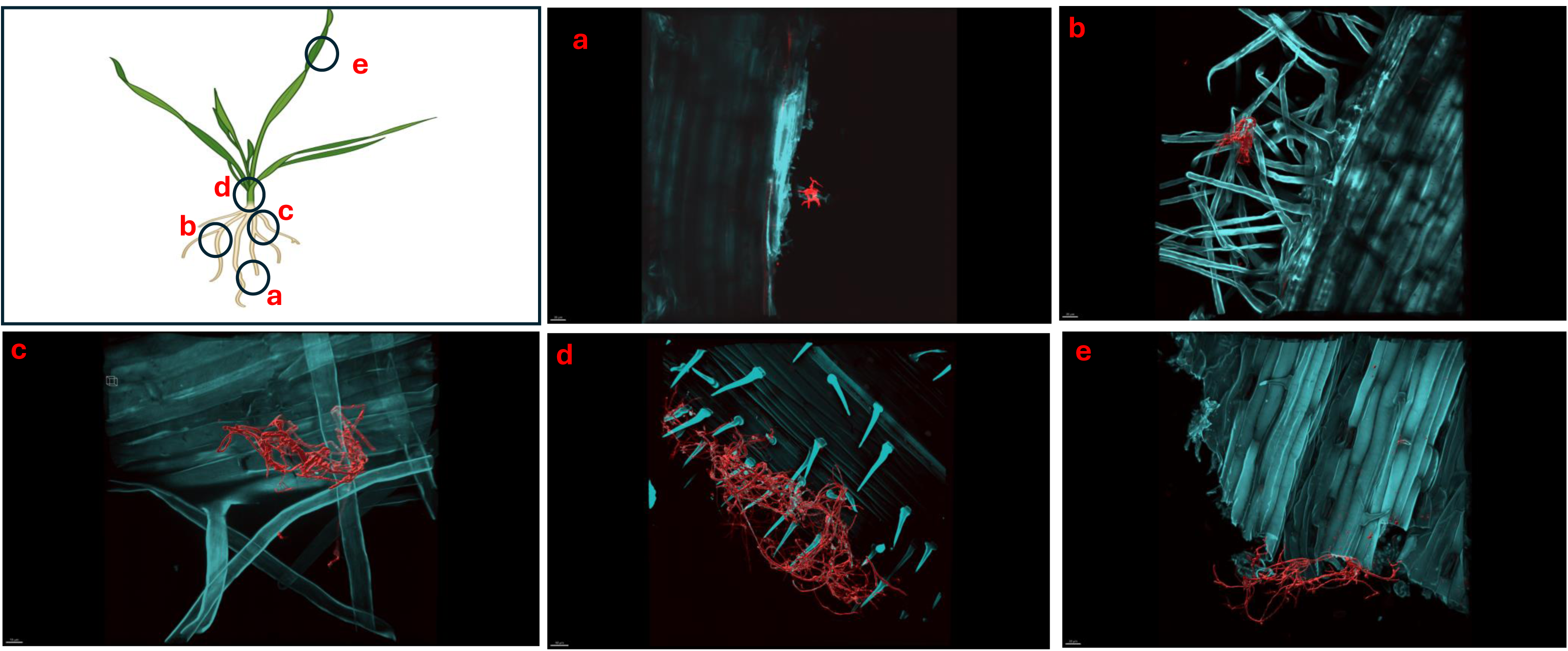
Micrographs showing *Triticum aestivum* plants inoculated with *Streptomyces coelicolor* M145 post-5-days of treatment. Samples were stained with both calcofluor white and wheat-germ agglutinin (WGA) 594. *Streptomyces coelicolor* M145 is visualized in red. The letters on the schematic (left) shows the sampling regions for microscopy. (A) scale bar = 30 µM, (B) scale bar = 30 µM, (C) scale bar = 10 µM, (D) scale bar = 40 µM, (E) scale bar = 30 µM.

Like day 5, imaging of day 7 and day 10 plants showed the similar pattern of colonization by *S. coelicolor* on wheat plants (See Fig. 5-6). Visible colonization of *S. coelicolor* was observed across all five imaged regions. Beginning at the root tip region, the morphology of *S. coelicolor* appeared more advanced compared to day 5, revealing defined cross-wall formations extending from the vegetative hyphae with clear attachment to root hairs. (See Fig. 5-6). Further, the lateral root formation region (region b) imaged on day 7 exhibited extensive growth and advanced morphology of *S. coelicolor* with hyphae formation, demonstrating *S. coelicolor*’s strong attachment to the multiple areas of the root hairs. Further up the root in the crown region revealed a cluster of *S. coelicolor* like those observed in other regions of the root on day 7 post-inoculation. This image, taken further away from the root itself, highlights *S. coelicolor*’s capability to colonize root hairs beyond the immediate vicinity of the root.

**Figure 5:**
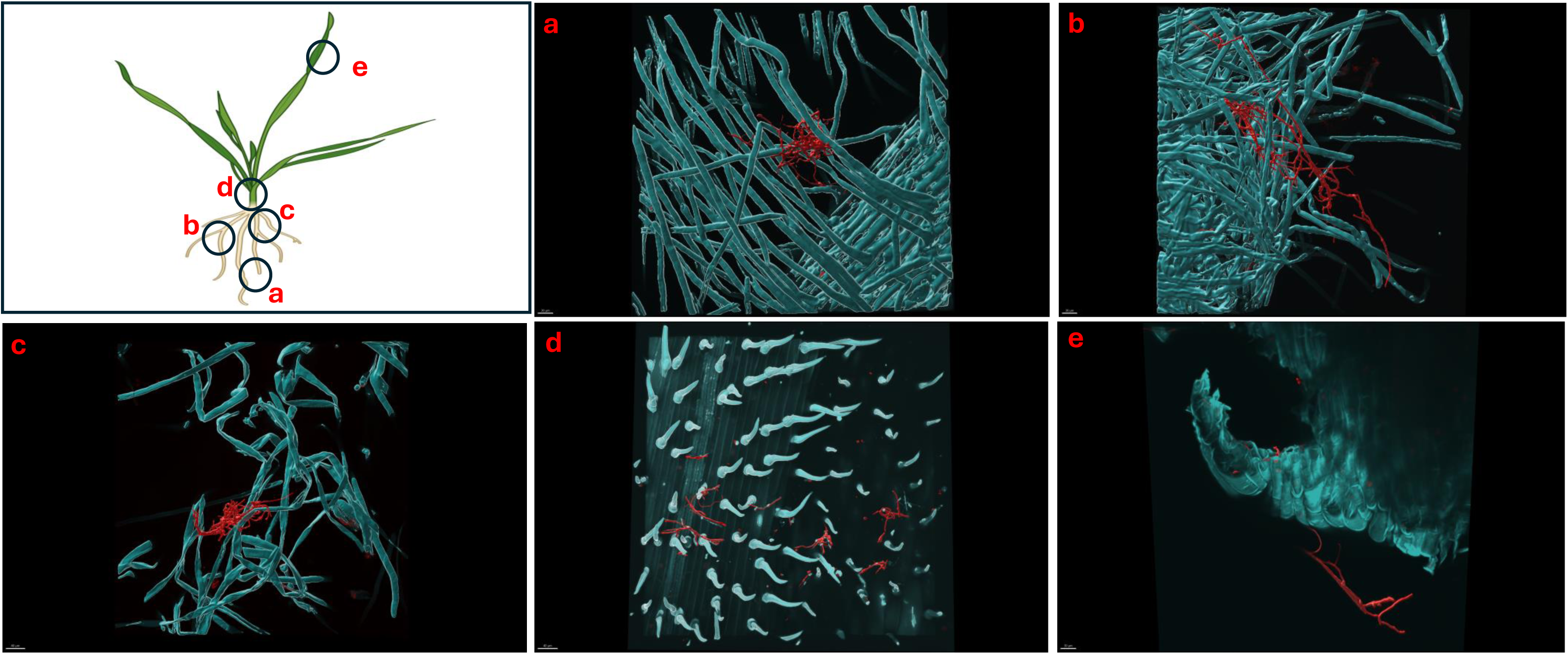
Micrographs showing *Triticum aestivum* plants inoculated with *Streptomyces coelicolor* M145 post-7-days of treatment. Samples were stained with both calcofluor white and wheat-germ agglutinin (WGA) 594. *Streptomyces coelicolor* M145 is visualized in red. The letters on the schematic (left) shows the sampling regions for microscopy. (A) scale bar = 30 µM, (B) scale bar = 30 µM, (C) scale bar = 40 µM, (D) scale bar = 40 µM, (E) scale bar = 30 µM.

Beyond the root region, the internode was examined for colonization. However, excessive pressure applied to the larger diameter of the internode impaired the full view of structure and morphology of *S. coelicolor*, making the analysis less interpretable. Despite this, the imaging revealed clear cross-wall formations, vegetative hyphae extensions, and evident colonization of *S. coelicolor* around the trichomes of the internodes. Finally, the shoot from day 7 post-inoculation was analyzed and revealed less abundant S. *coelicolor* colonizing the *T. aestivum*, positioned at a distance from the shoot. Lastly, day 10 *T. aestivum* plants were harvested and imaged (Fig. 6). At the root tip region, a substantial colony of *S. coelicolor* was identified, with the bacteria visibly wrapping around the entire left side of the root tip, suggesting that *S. coelicolor* can colonize the root tip itself. Moving beyond the root tip, the middle root image (region b) showed *S. coelicolor* extensively wrapping around a single root hair, encompassing it from all angles. Within this single *S. coelicolor* colony, two sporulating regions were evident (see Fig. 6). Continuing from the middle root, filamentous *S. coelicolor* were observed intertwining through the root hairs. Although this region corresponds to the mature root area, there appear to be fewer distinct cross-wall formations. Further beyond the root region, the internode exhibited the most extensive hyphae formations observed in the day 10 analysis. This region displays dense hyphae formations surrounding the trichomes of the internode. Notably, the hyphae curve around the base of the trichomes in multiple areas, which are potential entry points into the wheat plant for ingression. If *S. coelicolor* exhibits endophytic behavior in this in-vitro analysis, it is possible that the bacteria are gravitating toward these openings as a mechanism for plant entry. To verify our hypothesis that *S. coelicolor* may harness wheat plants to adapt to endophytic style, we conducted experiments wherein wheat plants were inoculated with *S. coelicolor* at the root level. Leaf blades were harvested post days 5/7/10 days of *S. coelicolor* inoculation. The cut leaf blades were surface sterilized with 50% ethanol for 10 seconds and placed on rich media. Our data showed that the leaf blades post surface sterilization led to formation and establishment of *S. coelicolor* colonies once plated (SOM Fig. 4). This data shows two distinct explanations, one that *S. coelicolor* moves extensively on the plant surface to reach the aerial parts or *S. coelicolor* may adapt to endophytic lifestyle on wheat plants.

**Figure 6:**
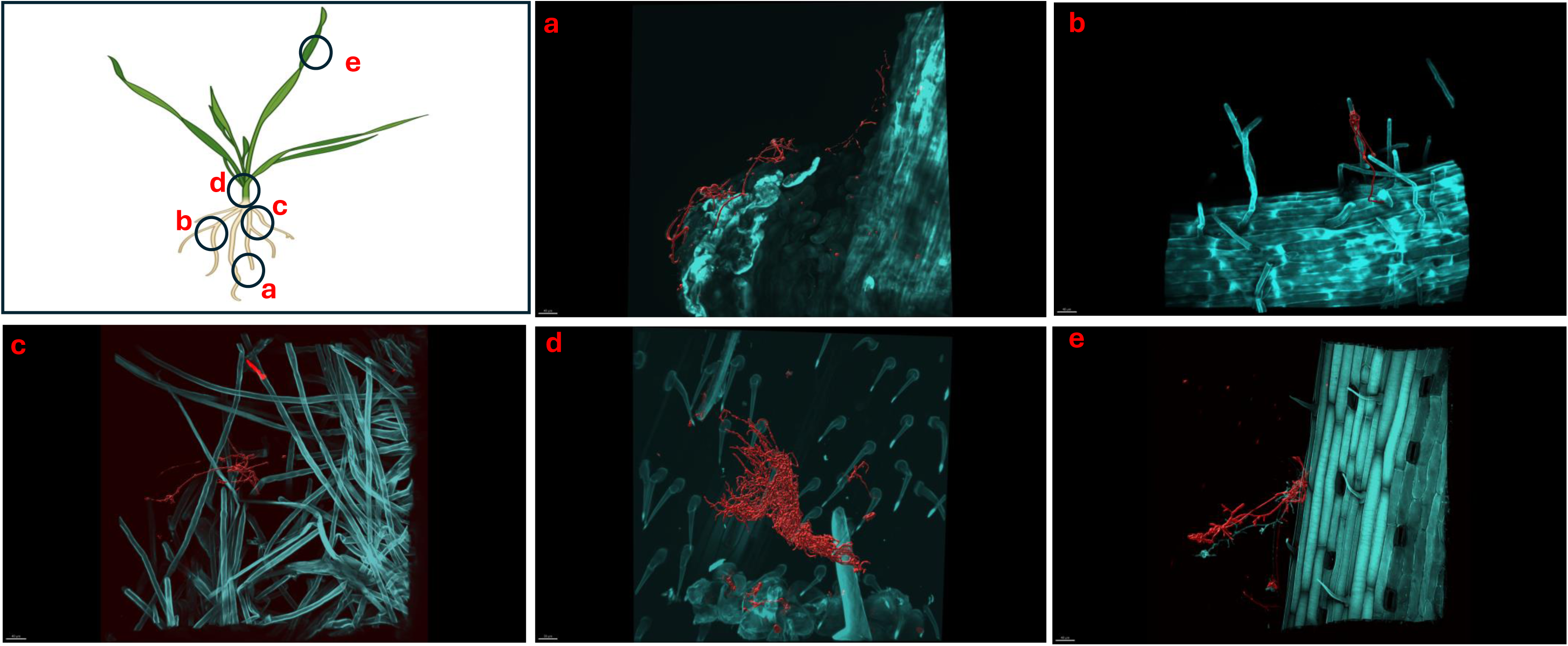
Micrographs showing *Triticum aestivum* plants inoculated with *Streptomyces coelicolor* M145 post-10-days of treatment. Samples were stained with both calcofluor white and wheat-germ agglutinin (WGA) 594. *Streptomyces coelicolor* M145 is visualized in red. The letters on the schematic (left) shows the sampling regions for microscopy. (A) scale bar = 40 µM, (B) scale bar = 40 µM, (C) scale bar = 40 µM, (D) scale bar = 30 µM, (E) scale bar = 40 µM.

### Extraction of EGT from *Triticum aestivum* roots and shoots post *Streptomyces coelicolor* M145 inoculation

*Triticum aestivum* plants inoculated with *S. coelicolor* M145 along with control plants were harvested on days 5 and 10, and EGT was detected and quantified from the roots and shoots of *T. aestivum* [SOM Fig. 1, Fig. 7A]. By day 5, quantifiable amounts of EGT were detectable in both roots and shoots. On the day 5 roots, trace amounts of EGT were recovered, alongside an increase in EGT on day 10 roots. Statistical analysis revealed the increase was present but not statistically significant.

**Figure 7:**
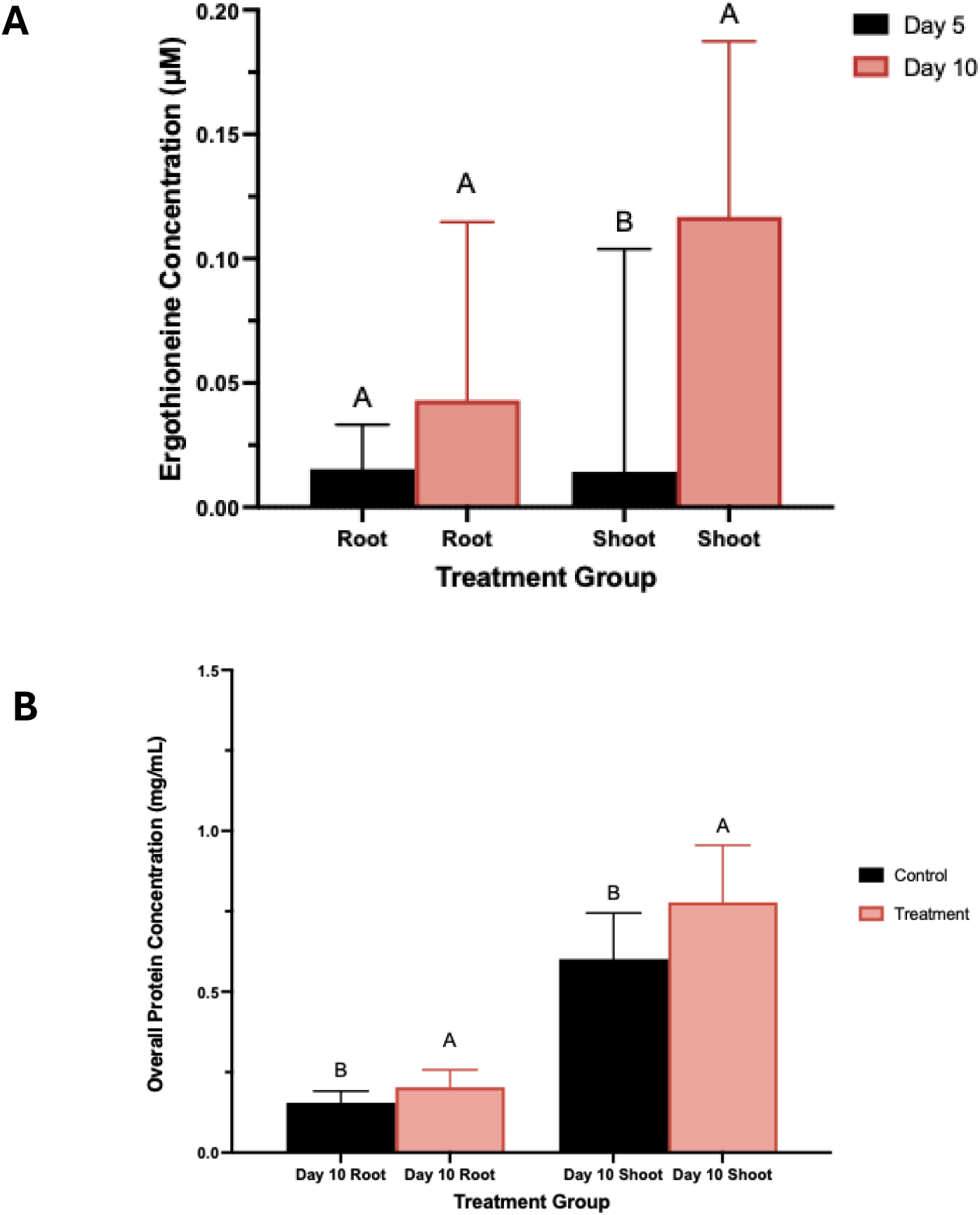
A) Total Ergothioneine concentration in wheat plants treated with *S. coelicolor*. Plants were treated with *S. coelicolor* and harvested on day 5 and 10. Roots and shoots were extracted for the total EGT concentration post bacterial treatment. Letters indicate statistically significant differences (p < 0.05) between treatments. B) Total protein concentration in wheat plants treated with *S. coelicolor* and harvested on day 5 and 10. Roots and shoots were extracted for the total protein concentration post bacterial treatment. Letters indicate statistically significant differences (p< 0.05) between treatments.

Meanwhile, the EGT extract from the shoot data showed a slightly different trend, with day 5 shoots containing trace amounts of EGT. By day 10, the shoots had accumulated significantly more EGT, overall indicating that EGT accumulates in a time-dependent manner in both roots and shoots of *Triticum aestivum* plants inoculated with *S. coelicolor*.

### Inoculation of *Streptomyces coelicolor* M145 on *Triticum aestivum* significantly enhances overall protein concentration

In the control *T. aestivum* roots, a lower protein concentration was detected compared to the *S. coelicolor*-treated plants. This result was significant, indicating that treatment of *S. coelicolor* on *T. aestivum* in-vitro leads to an increase in total protein concentration in both aboveground and belowground plant organs (see Fig 7B). Comparatively, the control *T. aestivum* roots/shoots exhibited a lower protein concentration than the *S. coelicolor*-treated plants.

## Discussion

### EGT biosynthesis declines under nutrient stress conditions

Actinobacteria, cyanobacteria, and some Methylobacteria species are known to biosynthesize EGT^27^. Additionally, some fungal species, including *Neurospora crassa*, *Cordyceps militaris*, and mushroom fruiting bodies, also biosynthesize EGT^24^. A previous study found that EGT levels were quantifiable in both intracellular and extracellular compartments of Mycobacterium smegmatis under normal growth conditions, with significantly higher extracellular EGT titers, suggesting active secretion of EGT^28^. Alternatively, in our study, we focused on quantifying intracellular EGT in *S. coelicolor*, where we hypothesized that growth media conditions play a role in *S. coelicolor* morphology and EGT production. The goal of our experiment was to compare intracellular EGT titers in *S. coelicolor* grown on its optimal medium, to those in a significantly less nutrient-rich medium formulated for plant health. Our study found that, at all three time points of extract harvest, EGT levels were consistently higher when was grown on a more nutrient rich media. We argue, that EGT being a secondary metabolite, its production is likely favored when primary metabolic needs are sufficiently met. To elaborate, primary metabolites are essential for cellular growth and development, and typically include amino acids, organic acids, and carbohydrates^29^. When the primary metabolites are sufficiently produced, the organisms may redirect its resources towards production of secondary metabolites, such as EGT. This indicates that EGT is not involved in the essential survival of *S. coelicolor*, but can present as a stress protector, ultimately providing a competitive ecological advantage over other organisms^29^. These findings also prove *S. coelicolor*’s metabolic robustness in the production of EGT, making large-scale fermentation processes of this amino acid more effective, resourceful, and potentially more cost productive. The implications of EGT for improving human health, coupled with its role as a secondary metabolite in *S. coelicolor*, suggest that this amino acid could be a promising candidate for utilization in agrochemical and pharmacological applications.

*S. coelicolor*-plant association and *in planta* EGT production.

Previous literature has shown that in soil experiments, low levels of EGT can be detected in plants due to the presence of EGT-producing fungi or bacteria^22^. Specifically, low levels of EGT have been found in plants such as garlic, wheat, oats, and beans when grown in association with EGT- producing microorganisms^22^. It is assumed that these microorganisms attach to the plant roots and often function as endophytes, entering the roots and subsequently producing EGT within the plant^30^. Amongst the Actinobacteria class, a prior study utilized scanning electron microscopy (SEM) to isolate and identify 482 *Streptomyces* strains as endophytes from twenty-eight different plant species. Specifically in spring wheat (*Triticum aestivum*), nine different native *Streptomyces* strains were identified, confirming the idea that *Streptomyces* may act as root endophyte^16^. Also, z-stacking has been previously implemented in research to study the relationship between motile and non-motile endophytic bacteria and their colonization patterns in *Musa acuminata* species^31^. Using this method, the authors observed a high abundance of live bacteria along the periphery of host cells, specifically within the peri-space between the cell wall and plasma membrane^31^. In our study, we hypothesized that *S. coelicolor* colonization to *T. aestivum* and potential ingression by bacteria may show in planta EGT production. As a result, we developed a fluorescent confocal microscopy protocol to analyze *S. coelicolor* colonization and its potential as an endophyte in spring wheat under *in-vitro* conditions. Upon successfully developing a fixation and staining protocol, we determined that *S. coelicolor* utilizes its hyphal extension properties in combination with plant root interactions to distribute itself throughout the three root regions imaged: root tip, middle root, and upper root (maturation zone). In addition to root colonization, we observed that *S. coelicolor* also colonizes the nodal plane and leaf blade regions. Interestingly, in the shoot region, *S. coelicolor* was detected interlacing at the site where the shoot was cut for microscopy preparation. This distribution of *S. coelicolor* to the internodes and shoots can be justified by a variety of reasoning. Since *S. coelicolor* is nonmotile, it typically relies on water, wind, or other organisms for movement^32^. One possible explanation for the microscopy analysis from the *in-vitro* experiment is that, due to the plant being sealed in the system with a liquid bacterial inoculum, along with consistent transpiration and temperature fluctuations between the petri plate and the external environment, the bacteria may have transferred themselves via water to the shoot regions, moving away from their initial inoculation site at the roots. Alternatively, *S. coelicolor* may have found an entry point and exhibited endophytic behavior, which could explain its presence in the area where the shoot was excised.

Further analysis using LC-MS/MS revealed quantifiable trace levels of EGT in both the roots and shoots of the plant following the inoculation of *S. coelicolor* on *T. aestivum*. By day 10 post-inoculation of bacteria, both the roots and shoots exhibited significantly higher levels of EGT. Similarly, by day 10, fresh weight measurements of both roots and shoots were significantly higher in treated plants compared to controls. These findings confirm our hypothesis that *S. coelicolor* contributes to EGT enhancement of *T. aestivum*. Beyond enhancing EGT levels in planta, we aimed to determine whether protein concentration increased in *T. aestivum* inoculated with *S. coelicolor.* A previous study confirmed that inoculation with three different PGPR’s under greenhouse conditions increased overall protein concentration in *Cucumis sativus L*. fruit with respect to the control^33^. In our study, *S. coelicolor* inoculation of *T. aestivum* in-vitro significantly enhanced overall protein concentration in both roots and shoots, suggesting that *S. coelicolor* plays a role in enhancing nutrient composition, presumably through its PGPR associated mechanisms including improved nutrient uptake or metabolic regulation.

## Conclusions

Taken together, our research demonstrates that the biosynthesis of EGT by *S. coelicolor* suggests its significant role in microbial stress responses, while also highlighting its biotechnological potential. Using the LC-MS/MS method we developed to detect EGT extracted from *S. coelicolor* cells, we found that EGT is biosynthesized as a secondary metabolite at trace levels under both nutrient rich and minimal conditions. Our findings also suggest that *S. oelicolor* inoculation in plants to enhance EGT and overall protein concentration, which may be beneficial to fortify cereal plants for enhanced protein concentration in agriculture. Through our development of a confocal fluorescent microscopy protocol, inoculation of *S. coelicolor* on *T. aestivum* resulted in visible colonization of roots, shoots, and internodes. This colonization corresponded with enhanced EGT and biomass levels in both root and shoots. These findings suggest that *S. coelicolor* functions endophytically, contributing to EGT biosynthesis and promoting plant growth and nutrient accumulation within the plant. Given the growing relevance of EGT in both agricultural and medical contexts, this research further establishes *S. coelicolor* as a valuable organism for future studies on its potential applications for both plant and human physiological studies. In addition, the role of EGT biosynthesizing soil microbes for plant association opens a new discussion to evaluate the soil to human health continuum in the context of crop fortification by harnessing soil/plant microbiome.

## Supporting information

SOM Figures 1-4

## Acknowledgements

The work was conducted with the grant received by authors from USDA FFAR to JK, AS, GZ and HB. Authors acknowledge the support and help from Mathew Traxler’s group at University of California, Berkeley for culturing and maintaining *Streptomyces coelicolor* M145.

## Legends to supplementary figures

SOM Figure 1: Resulting mass spectrometry data from the TOF-MS/MS profile of a pure ergothioneine standard (a); *S. coelicolor* cell extracts (b) Intracellular *S. coelicolor* M145 intracellular extracts and (c) wheat root extracts.

SOM Figure 2: Shoot fresh weight (g) of *Triticum aestivum* plants in control and treatment groups post-inoculation with *Streptomyces coelicolor* M145. The panels (A-B) show morphological traits in control and *S. coelicolor* inoculated plants on day 5 and 10. Asterisk indicates statistically significant differences (p < 0.05) between treatments and control.

SOM Figure 3: Microscopy of *Triticum aestivum* plants untreated with *Streptomyces coelicolor* M145. The images were taken post 10 days of growth cycle. Samples were stained with both calcofluor white and wheat-germ agglutinin (WGA) 594. The letters on the schematic (left) shows the sampling regions for microscopy. (A) scale bar = 40 µM, (B) scale bar = 40 µM, (C) scale bar = 40 µM, (D) scale bar = 40 µM, (E) scale bar = 40 µM.

SOM Figure 4: Protocol developed to evaluate the endophytic lifestyle of *S. coelicolor* in wheat plants. Wheat plants post root inoculation with *S. coelicolor* were harvested on day 5, 7 and 10, leaf blades from the harvested plants were surface sterilized using 50% ethanol for 15 seconds and were plated on a rich media for *S. coelicolor* colony formations. The control untreated sample on the left depicts plants not treated with *S. coelicolor* and detached leaves were plated on day 10 post culture.

## References

1. El-Ramady H, Hajdú P, Törős G, et al. Plant nutrition for human health: A pictorial review on plant bioactive compounds for sustainable agriculture. Sustainability. 2022;14(14):8329.

2. Huang S, Wang P, Yamaji N, Ma JF. Plant nutrition for human nutrition: Hints from rice research and future perspectives. Molecular Plant. 2020;13(6):825–835.

3. Priya AK, Alagumalai A, Balaji D, Song H. Bio-based agricultural products: A sustainable alternative to agrochemicals for promoting a circular economy. RSC Sustainability. 2023;1(4):746–762.

4. Shayanthan A, Ordoñez PAC, Oresnik IJ. The role of synthetic microbial communities (SynCom) in sustainable agriculture. Frontiers in Agronomy. 2022; 4:896307.

5. Basheer S, Wang X, Farooque AA, Nawaz RA, Pang T, Neokye EO. A review of greenhouse gas emissions from agricultural soil. Sustainability. 2024;16(11):4789.

6. Dhankhar N, Kumar J. Impact of increasing pesticides and fertilizers on human health: A review. Materials Today: Proceedings. 2023: S2214785323018382.

7. Silva V, Yang X, Fleskens L, Ritsema CJ, Geissen V. Environmental and human health at risk – scenarios to achieve the farm to fork 50\% pesticide reduction goals. Environ Int. 2022; 165:107296.

8. Miranda AM, Hernandez-Tenorio F, Villalta F, Vargas GJ, Sáez AA. Advances in the development of biofertilizers and biostimulants from microalgae. Biology. 2024;13(3):199.

9. Du Jardin P. Plant biostimulants: Definition concept, main categories and regulation. Scientia Horticulturae. 2015; 196:3–14.

10. Zhang C. Efficacy of oral administration of ergothioneine (DR. ERGO™) on skin health improvement: A single-center open-label clinical study. American Journal of Biomedical Science \& Research. 2023;20(6):822–828.

11. Ali O, Ramsubhag A, Jayaraman J. Biostimulant properties of seaweed extracts in plants: Implications towards sustainable crop production. Plants. 2021;10(3):531.

12. Rengasamy, Sathya, Ushadevi, T. Industrially important enzymes producing streptomyces species from mangrove sediments. International Journal of Pharmacy and Pharmaceutical Sciences (IJPPS). 2014;6(10).

13. Viaene T, Langendries S, Beirinckx S, Maes M, Goormachtig S. *Streptomyces* as a plant’s best friend? FEMS Microbiol Ecol. 2016;92(8): fw119.

14. Nazari MT, Schommer VA, Braun JCA, et al. Using streptomyces spp. as plant growth promoters and biocontrol agents. Rhizosphere. 2023; 27:100741.

15. Vurukonda SSKP, Mandrioli M, D’Apice G, Stefani E. Draft genome sequence of plant growth- promoting Streptomyces sp. strain SA51 isolated from olive trees. Microbiology Resource Announcements. 2020;9(1):768.

16. Sardi P, Saracchi M, Quaroni S, Petrolini B, Borgonovi GE, Merli S. Isolation of endophytic Streptomyces strains from surface-sterilized roots. Appl Environ Microbiol. 1992;58(8):2691–2693.

17. Styczen ME, Abrahamsen P, Hansen S, Knudsen L. Model analysis of the significant drop in protein content in danish grain crops from 1990-2015. Eur J Agron. 2020; 118:126068.

18. Medek DE, Schwartz J, Myers SS. Estimated effects of future atmospheric CO2 concentrations on protein intake and the risk of protein deficiency by country and region. Environ Health Perspect. 2017;125(8):087002.

19. Seltenrich N. Estimated deficiencies resulting from reduced protein content of staple foods: Taking the cream out of the crop? Environ Health Perspect. 2017;125(9):094001.

20. Borodina I, Kenny LC, McCarthy CM, et al. The biology of ergothioneine, an antioxidant nutraceutical. Nutrition Research Reviews. 2020;33(2):190–217.

21. Smith E, Ottosson F, Hellstrand S, et al. Ergothioneine is associated with reduced mortality and decreased risk of cardiovascular disease. Heart. 2020;106(9):691–697.

22. Tian X, Thorne JL, Moore JB. Ergothioneine: An underrecognised dietary micronutrient required for healthy ageing? Br J Nutr. 2023;129(1):104–114.

23. Carrara JE, Lehotay SJ, Lightfield AR, et al. Linking soil health to human health: Arbuscular mycorrhizae play a key role in plant uptake of the antioxidant ergothioneine from soils. *PLANTS, PEOPLE*, PLANET. 2023;5(3):449–458.

24. Chen Z, He Y, Wu X, Wang L, Dong Z, Chen X. Toward more efficient ergothioneine production using the fungal ergothioneine biosynthetic pathway. Microbial Cell Factories. 2022;21(1):76.

25. Nakajima S, Satoh Y, Yanashima K, Matsui T, Dairi T. Ergothioneine protects streptomyces coelicolor A3(2) from oxidative stresses. Journal of Bioscience and Bioengineering. 2015;120(3):294– 298.

26. Murashige T, Skoog F. A revised medium for rapid growth and bioassays with tobacco tissue cultures. Physiol Plantarum. 1962;15(3):473–497.

27. Cumming BM, Chinta KC, Reddy VP, Steyn AJC. Role of ergothioneine in microbial physiology and pathogenesis. Antioxidants \& Redox Signaling. 2018;28(6):431–444.

28. Sao Emani C, Williams MJ, Wiid IJ, et al. Ergothioneine is a secreted antioxidant in mycobacterium smegmatis. Antimicrob Agents Chemother. 2013;57(7):3202–3207.

29. Lee D, Yoon MH, Kang YP, et al. Comparison of primary and secondary metabolites for suitability to discriminate the origins of schisandra chinensis by GC/MS and LC/MS. Food Chem. 2013;141(4):3931– 3937.

30. Eung-Jun Park*, Wi Young Lee, Seung Taek Kim, Jin Kwon Ahn and Eun Kyung Bae. Ergothioneine accumulation in a medicinal plant gastrodia elata. Journal of Medicinal Plants Research. 2010;4(12).

31. Thomas P, Reddy KM. Microscopic elucidation of abundant endophytic bacteria colonizing the cell wall–plasma membrane peri-space in the shoot-tip tissue of banana. AoB PLANTS. 2013;5.

32. Schlimpert S, Elliot MA. The best of both Worlds—Streptomyces coelicolor and streptomyces venezuelae as model species for studying antibiotic production and bacterial multicellular development. J Bacteriol. 2023;205(7):153.

33. Bhardwaj RL, Parashar A, Parewa HP, Vyas L. An alarming decline in the nutritional quality of foods: The biggest challenge for future generations’ health. Foods. 2024;13(6):877.

34. Zapata-Sifuentes G, Hernandez-Montiel LG, Saenz-Mata J, et al. Plant growth-promoting rhizobacteria improve growth and fruit quality of cucumber under greenhouse conditions. Plants. 2022;11(12):1612.

35. Alam K, Mazumder A, Sikdar S, et al. Streptomyces: The biofactory of secondary metabolites. Frontiers in Microbiology. 2022; 13:968053.

36. 36. Halpern M, Bar-Tal A, Ofek M, Minz D, Muller T, Yermiyahu U. The use of biostimulants for enhancing nutrient uptake. In: Advances in {agronomy}. Vol 130. Elsevier; 2015:141–174.

37. Kartseva T, Alqudah AM, Aleksandrov V, et al. Nutritional genomic approach for improving grain protein content in wheat. Foods. 2023;12(7):1399

38. Mena P, Angelino D. Plant food nutrition and human health. Nutrients. 2020;12(7):2157.

34. Bhardwaj RL, Parashar A, Parewa HP, Vyas L. An alarming decline in the nutritional quality of foods: The biggest challenge for future generations’ health. Foods. 2024;13(6):877.

36. Chow YW, Pietranico R, Mukerji A. Studies of oxygen binding energy to hemoglobin molecule. Biochem Biophys Res Commun. 1975;66(4):1424–1431.

42. 37. Halpern M, Bar-Tal A, Ofek M, Minz D, Muller T, Yermiyahu U. The use of biostimulants for enhancing nutrient uptake. In: Advances in {agronomy}. Vol 130. Elsevier; 2015:141–174.

38. Kartseva T, Alqudah AM, Aleksandrov V, et al. Nutritional genomic approach for improving grain protein content in wheat. Foods. 2023;12(7):1399.

39. Mena P, Angelino D. Plant food nutrition and human health. Nutrients. 2020;12(7):2157.

40. Osborn D, Jenkinson DH. Proceedings: Comparison of the effects of selective alpha and beta-receptor agonists on intracellular cyclic AMP levels and glycogen phosphorylase activity in guinea-pig liver. Br J Pharmacol. 1975;55(2):286P–287P.

41. Schmoldt A, Benthe HF, Haberland G. Digitoxin metabolism by rat liver microsomes. Biochem Pharmacol. 1975;24(17):1639–1641.

42. Shi X, Ling H. Current advances in genome sequencing of common wheat and its ancestral species. The Crop Journal. 2018;6(1):15–21.

43. Schmoldt A, Benthe HF, Haberland G. Digitoxin metabolism by rat liver microsomes. Biochem Pharmacol. 1975;24(17):1639–1641.

44. Sun W, Shahrajabian MH, Soleymani A. The roles of plant-growth-promoting rhizobacteria (PGPR)- based biostimulants for agricultural production systems. Plants. 2024;13(5):613.

45. Tarmure S, Alexescu T, Orasan O, et al. Influence of pesticides on respiratory pathology – a literature review. Annals of Agricultural and Environmental Medicine. 2020;27(2):194–200.

46. Tien Lea D, Duc Chua H, Quynh Lea N. Improving nutritional quality of plant proteins through genetic engineering. Curr Genomics. 2016;17(3):220–229.

47. Zapata-Sifuentes G, Hernandez-Montiel LG, Saenz-Mata J, et al. Plant growth-promoting rhizobacteria improve growth and fruit quality of cucumber under greenhouse conditions. Plants. 2022;11(12):1612

43. Shi X, Ling H. Current advances in genome sequencing of common wheat and its ancestral species. The Crop Journal. 2018;6(1):15–21

44. Sonnhof U, Grafe P, Krumnikl J, Linder M, Schindler L. Inhibitory postsynaptic actions of taurine, GABA and other amino acids on motoneurons of the isolated frog spinal cord. Brain Res. 1975;100(2):327–341

45. Sun W, Shahrajabian MH, Soleymani A. The roles of plant-growth-promoting rhizobacteria (PGPR)- based biostimulants for agricultural production systems. Plants. 2024;13(5):613.

46. Tarmure S, Alexescu T, Orasan O, et al. Influence of pesticides on respiratory pathology – a literature review. Annals of Agricultural and Environmental Medicine. 2020;27(2):194–200.

47. Tien Lea D, Duc Chua H, Quynh Lea N. Improving nutritional quality of plant proteins through genetic engineering. Curr Genomics. 2016;17(3):220–229.

