## Supplementary material for "*Streptomyces coelicolor*-plant association facilitates ergothioneine (EGT) uptake in *Triticum aestivum*": SOM Figures 1-4

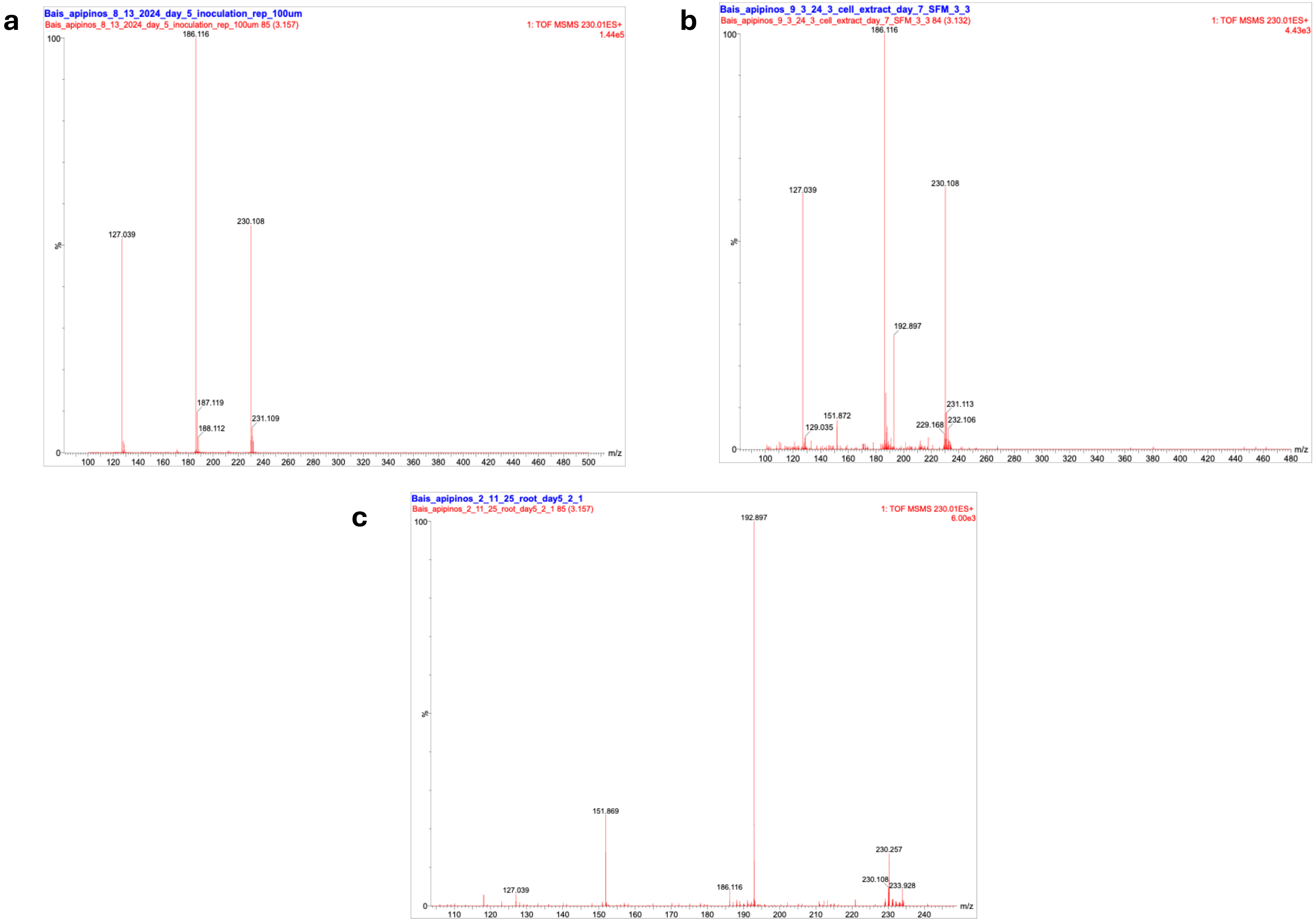


**SOM Figure 1** : Resulting mass spectrometry data from the TOF-MS/MS profile of a pure ergothioneine standard (a); *S. coelicolor* cell extracts (b) Intracellular *S. coelicolor* M145 intracellular extracts and (c) wheat root extracts.

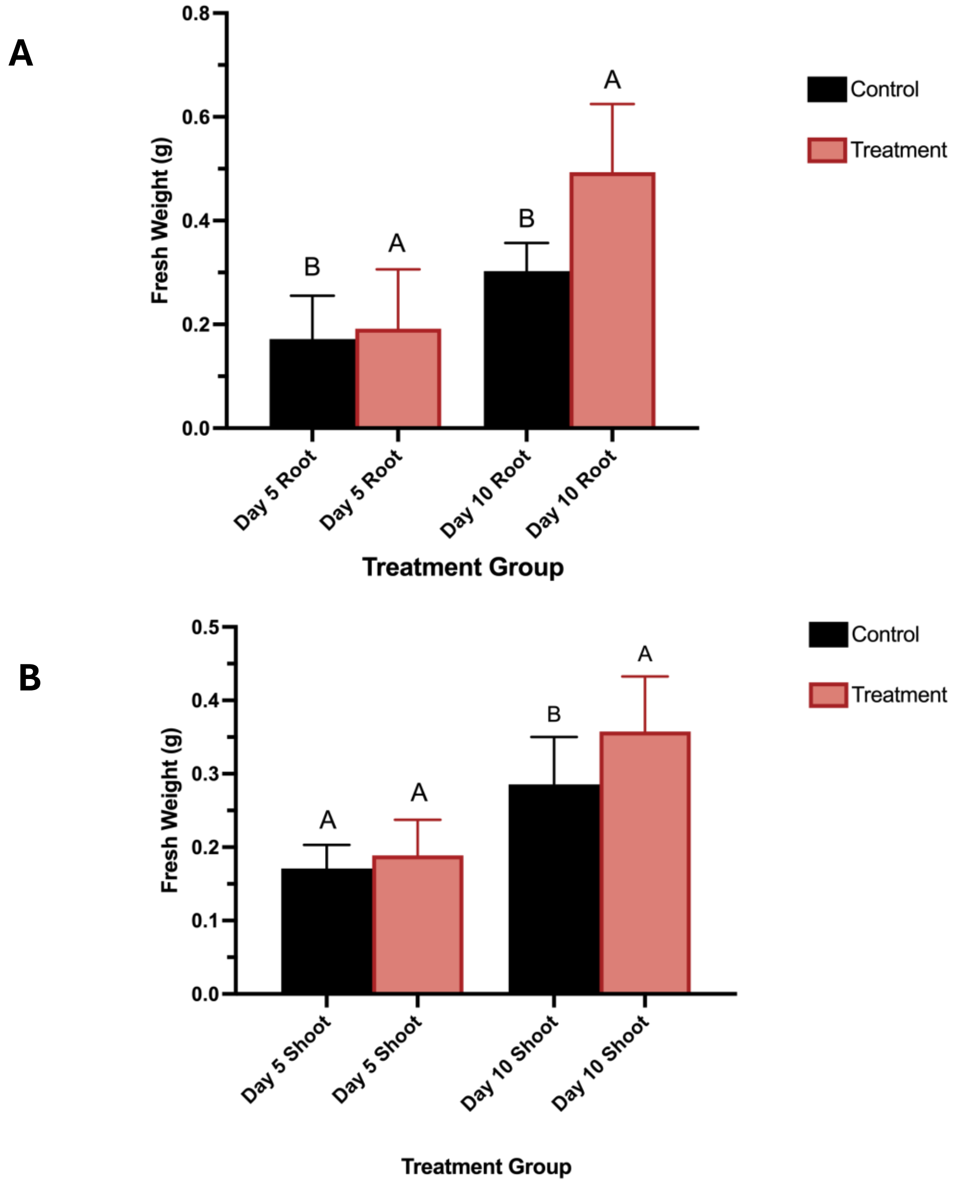


**SOM Figure 2:** Root and shoot fresh weight (g) of *Triticum aestivum* plants in control and treatment groups post-inoculation with *Streptomyces coelicolor* M145. The panels (A-B) shows morphological traits in control and *S. coelicolor* inoculated plants on day 5 and 10. Letters indicates statistically significant differences ($p < 0.05$) between treatments and control

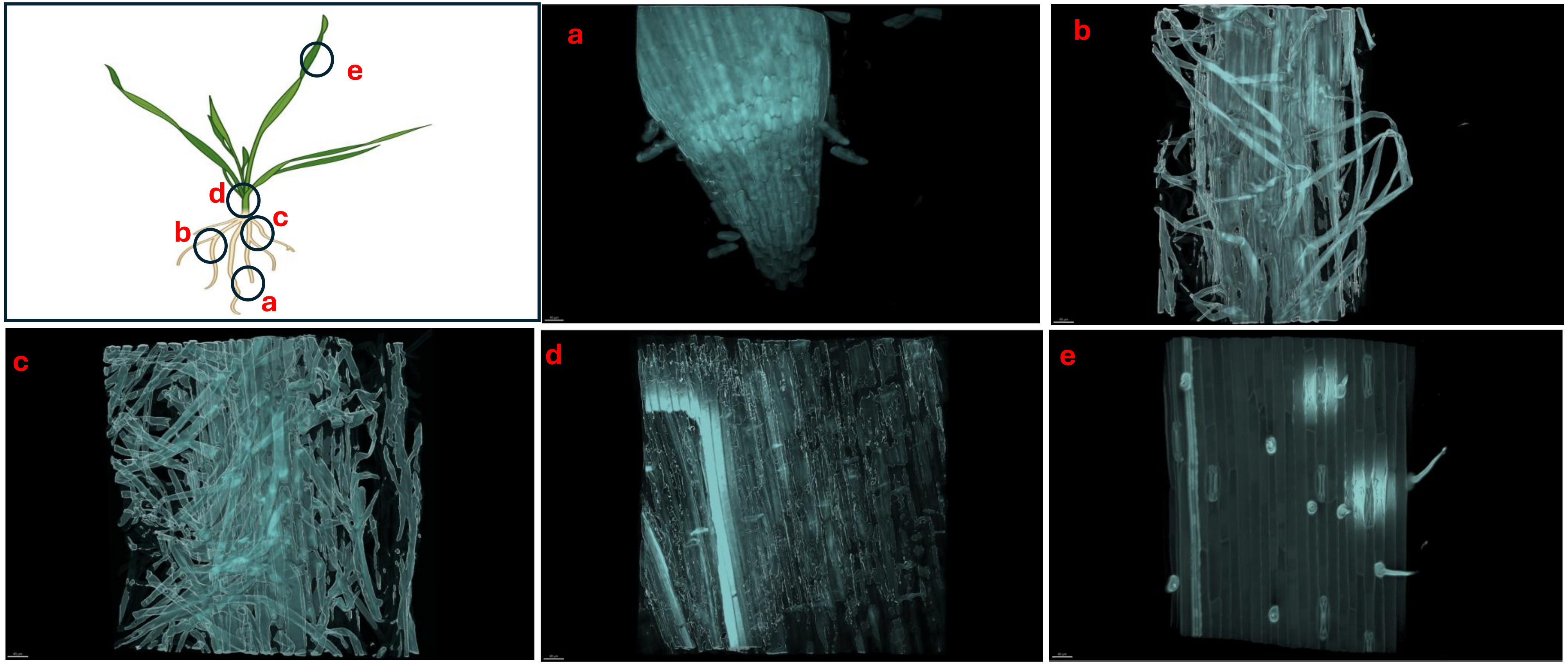


**SOM Figure 3:** Microscopy of *Triticum aestivum* plants untreated with *Streptomyces coelicolor* M145. The images were taken post 10 days of growth cycle. Samples were stained with both calcofluor white and wheat-germ agglutinin (WGA) 594. The letters on the schematic (left) shows the sampling regions for microscopy. (A) scale bar = 40 µM, (B) scale bar = 40 µM, (C) scale bar = 40 µM, (D) scale bar = 40 µM, (E) scale bar = 40 µM.

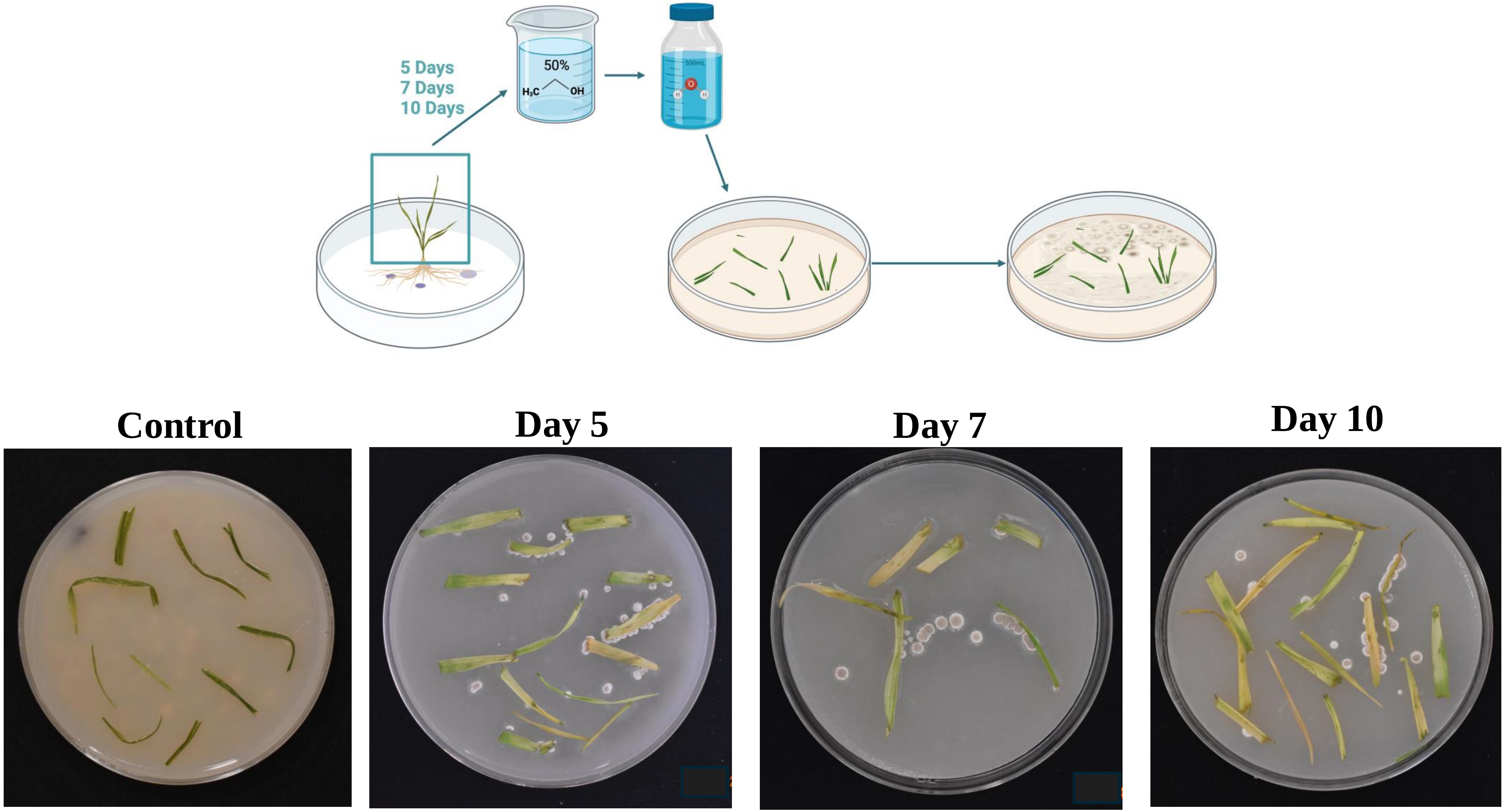


**SOM Figure 4 :** Protocol developed to evaluate the endophytic lifestyle of *S. coelicolor* in wheat plants. Wheat plants post root inoculation with *S. coelicolor* wee harvested on day 5, 7 and 10, leaf blades from the harvested plants were surface sterilized using 50% ethanol for 15 seconds and were plated on a rich media for *S. coelicolor* colony formations. The control untreated sample on the left depicts plants not treated with *S. coelicolor* and detached leaves were plated on day 10 post culture.
